# Early-life stress triggers long-lasting organismal resilience and longevity via tetraspanin

**DOI:** 10.1101/2023.07.25.550452

**Authors:** Wei I. Jiang, Henry De Belly, Bingying Wang, Andrew Wong, Minseo Kim, Fiona Oh, Jason DeGeorge, Xinya Huang, Shouhong Guang, Orion D. Weiner, Dengke K. Ma

## Abstract

Early-life stress experiences can produce lasting impacts on organismal adaptation and fitness. How transient stress elicits memory-like physiological effects is largely unknown. Here we show that early-life thermal stress strongly up-regulates *tsp-1*, a gene encoding the conserved transmembrane tetraspanin in *C. elegans*. TSP-1 forms prominent multimers and stable web- like structures critical for membrane barrier functions in adults and during aging. The up- regulation of TSP-1 persists even after transient early-life stress. Such regulation requires CBP- 1, a histone acetyl-transferase that facilitates initial *tsp-1* transcription. Tetraspanin webs form regular membrane structures and mediate resilience-promoting effects of early-life thermal stress. Gain-of-function TSP-1 confers marked *C. elegans* longevity extension and thermal resilience in human cells. Together, our results reveal a cellular mechanism by which early-life thermal stress produces long-lasting memory-like impact on organismal resilience and longevity.

**Teaser:** Studies reveal mechanisms of how early-life heat exposure produces long-lasting benefits on longevity in the nematode *C. elegans*.

## Introduction

Epidemiological and clinical studies in humans show that life stress of various forms can exert profound lasting impacts on mental and physical health outcomes and lifespans (*1–4*). For example, severe nutritional stress early in life can incur lifespan costs while psychosocial early- life stress increases vulnerability to psychiatric disorders (*5–7*). By contrast, milder physiological stresses, such as fasting with adequate nutrition or thermal stimuli via sauna exposure, are associated with long-lasting health benefits (*8, 9*). Transient periods of stress can induce persistent changes in the endocrine response, epigenetic regulation of gene expression and plasticity changes in various organs (*10–12*). However, the underlying molecular and cellular mechanisms by which transient early-life stress can produce memory-like physiological effects remain poorly understood. Additionally, it is challenging to establish causal relationship between putative cellular mechanisms and long-term health outcomes in human studies.

The free-living nematode *Caenorhabditis elegans* has emerged as a tractable model system to study how early-life stress may affect adult phenotypes. Early-life exposure to starvation, anoxia or osmotic stresses in *C. elegans* induces the transition into an L1 arrested developmental stage or stress-resistant dauer stage, which can persist until environmental conditions become favorable (*13–15*). Adults that have undergone the dauer stage preserve a memory of their early-life starvation experience, resulting in alterations in gene expression, extended lifespan, and decreased reproductive capacity (*16, 17*). Early-life starvation experience can also reorganize neuronal modulation of stage-specific behavioral responses to stresses (*18*). In addition, a one-day shift from 20 °C to 25 °C during early adulthood in *C. elegans* appears to improve stress resistance and extend lifespan through known stress- responding transcription factors: DAF-16, HSF-1, and HIF-1 (*19, 20*). It remains unclear how specific effectors of these transcription factors, or other epigenetic mechanisms independent of these factors, may elicit long-lasting impacts on adult stress resilience and longevity.

In this study, we use a robust thermal stress paradigm in *C. elegans* to uncover causal mechanisms by which transient stress may exert lasting impacts on organismal resilience and longevity. We show that transient heat exposure at 28 °C during late larval development activates the gene *tsp-1*, which encodes a *C. elegans* homolog of the evolutionarily conserved tetraspanin protein family. TSP-1 proteins form tetraspanin web-like structures and are essential for maintaining membrane permeability, barrier functions, and heat-induced organismal resilience and longevity. Initial induction of *tsp-1* by heat requires the histone acetyltransferase CBP-1; however, surprisingly, this results in sustained up-regulation of TSP-1 protein without a corresponding increase in mRNA abundance. We propose that the stability of the tetraspanin web-like structure, achieved through TSP-1 multimerization, mediates the long-lasting organismal impact triggered by transient mild thermal stress in early life.

## Results

Using RNA-seq, we previously identified *tsp-1* as one of the genes highly up-regulated by cold-warming (CW) stress (transient exposure to 4 or -20 °C followed by recovery at 20 °C) in *C. elegans* (*21–23*). We generated translational reporters by fusing GFP with endogenous regulatory DNA sequences (promoter and coding regions) for many of the CW stress-inducible genes to monitor their induction kinetics and protein localization in live *C. elegans* during development and adult responses to stresses. We focus on *tsp-1* in this study as the constructed *tsp-1*p::*tsp-1*::GFP translational reporter shows robust and striking induction at L4 stages by not only CW stresses, but also mild heat stress at 25 °C, more so at 28 °C, less so at 20 °C or above 32 °C (**Fig. 1A, B; fig. S1**). Induction by 28 °C is heat-duration dependent (**fig. S1**) and does not require HSF-1 (heat shock factor) (**fig. S2**) that mediates the canonical heat shock response in *C. elegans* (*24–28*). By contrast, the heat-shock chaperon gene *hsp-16* is

**Fig. 1.**
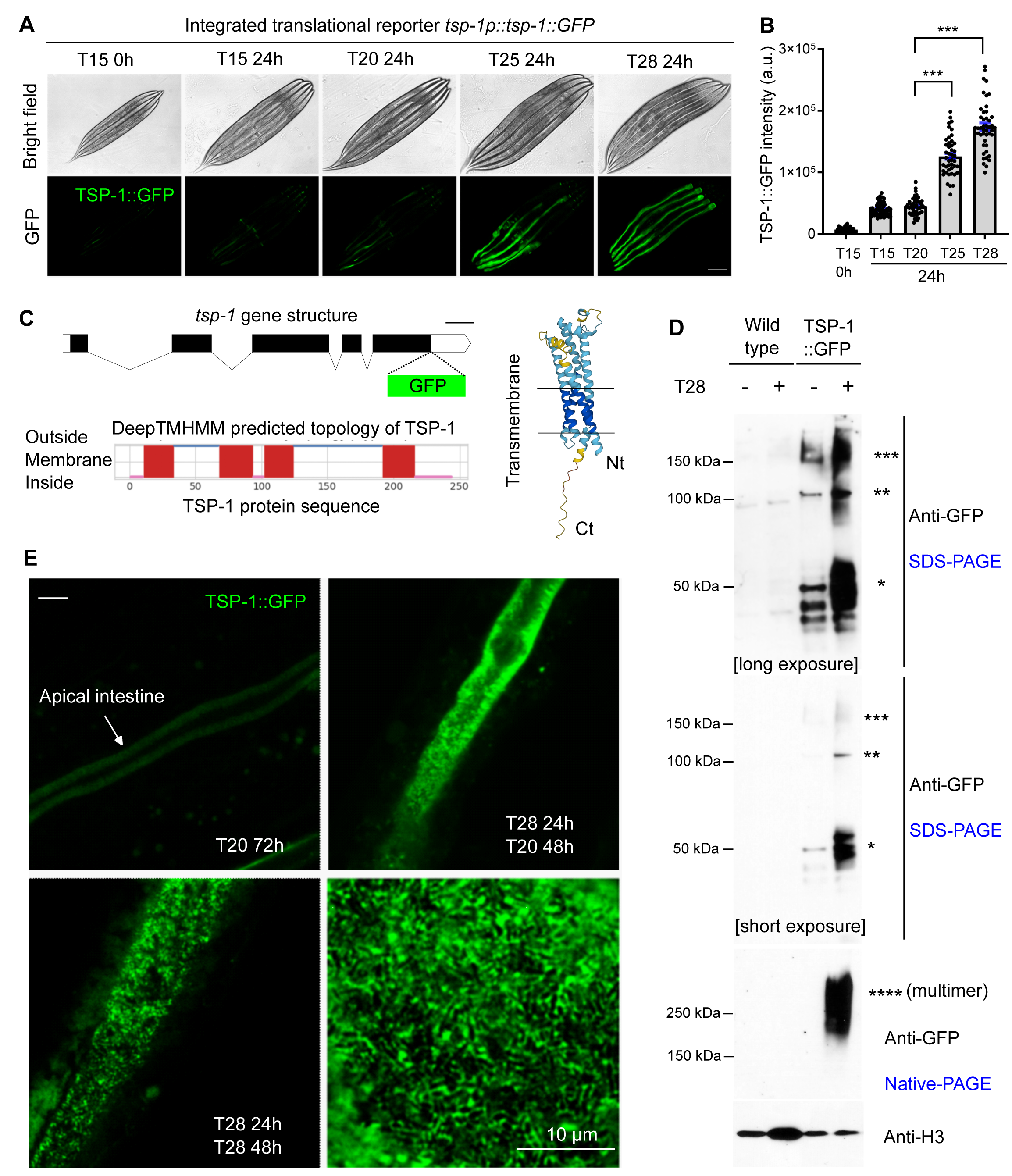
Thermal stress induces TSP-1 expression and tetraspanin web-like structure formation. (A) Representative bright field and epifluorescence images showing expression of an integrated transgene *tsp-1p::tsp-1::GFP* under temperatures and durations indicated. Scale bars: 100 µm. (B) Quantification of TSP-1::GFP fluorescence intensities under conditions indicated. *** indicates *P* < 0.001 (n > 40 animals per condition). (C) Schematic of the *tsp-1* gene structure, predicted membrane topology (by DeepTMHMM) and structure (by AlphaFold2) of TSP-1. (D) SDS-PAGE and native-PAGE western blots showing heat-induced increase in abundance and formation of dimers and multimers (asterisk) by TSP-1::GFP. (E) Representative confocal fluorescence images showing high-resolution Z-stack views of tetraspanin web structure formed by TSP-1::GFP, with enlarged view at right bottom. Scale bars: 10 µm.

also induced by 28 °C and requires HSF-1 for heat induction (**fig. S2**). These results highlight a previously unknown, HSF-independent thermal induction of TSP-1 in *C. elegans*.

TSP-1 is a *C. elegans* homolog of the evolutionarily conserved tetraspanin protein family in eukaryotes (*29–31*), predicted to form four-transmembrane α-helical segments and two extracellular loop domains (**Fig. 1C**). We fused GFP to the C-terminus of TSP-1, leaving the N-terminus of TSP-1 intact, which contains the first transmembrane helix targeting TSP-1 to plasma membranes (**Fig. 1C**). Tetraspanins are versatile scaffolding transmembrane proteins, specific members of which can interact with themselves to form multimers and control the spatial organization of membrane lipids and proteins in networks called tetraspanin webs or tetraspanin-enriched microdomains (*29–31*). We used sodium dodecyl-sulfate polyacrylamide gel electrophoresis (SDS-PAGE) and native PAGE to analyze the thermal induction and multimerization of TSP-1::GFP (**Fig. 1D**). Under baseline conditions at 20 °C, TSP-1::GFP formed protein species corresponding to predicted monomers, SDS-resistant dimers and trimers (in molecular weights). Heat at 28 °C for 24 hrs drastically increased all TSP-1::GFP species in SDS-PAGE, and native PAGE further revealed the formation of high molecular-weight species larger than dimers and trimers (**Fig. 1D**). Confocal microscopy revealed that TSP-1::GFP was highly enriched along the apical membrane of intestinal cells (**Fig. 1E**). Heat at 28 °C for 72 hrs or a transient period of 24 hrs starting from L4 markedly increased the abundance of TSP- 1::GFP, forming tetraspanin web-like structures discernable at high resolution (**Fig. 1E**). Such structures were not caused by temperature effect on the tag GFP *per se* since intestinal apical membrane-tethered GFP alone did not show such a pattern (**fig. S3**). In addition, *tsp-1* endogenously tagged with a worm-codon optimized fluorescent protein wrmScarlet by CRISPR/Cas showed similar up-regulation by heat at 28 °C and formation of tetraspanin web-like structures (**fig. S4**). Compared with multi-copy transgenic TSP-1::GFP, endogenously tagged TSP-1::wrmScarlet exhibited overall weaker fluorescent intensity and enabled us to better resolve finer tetraspanin webs with regular lattice-like structures (**fig. S4F-I**).

Tetraspanin has been implicated in numerous biological processes, yet its precise cellular functions remain largely unknown. We next sought to determine normal TSP-1 expression pattern under non-stress conditions and its adult physiological function during aging in *C. elegans*. At day 1 (young adult), 5 (adult), and 9 (old adult) post L4 stages, TSP-1::GFP exhibited a progressive increase in abundance (**Fig. 2A-E**). Cultivation at 25 °C accelerated the time-dependent increase of TSP-1::GFP abundance, albeit to a lesser degree than a transient 24 hrs period at 28 °C (**Fig. 2A-E**, **Fig. 1E**) (28 °C represents harsh stress precluding chronic analyses of TSP-1::GFP at day 5 and 9 because of organismal death by prolonged exposure, see below). The striking tetraspanin web-like structure formed by TSP-1::GFP (**Fig. 1E**) prompted us to test the role of TSP-1 in maintaining membrane integrity. We developed a fluorescein-based assay modified from our previous studies (*21*) to measure the acute barrier function or permeability of intestinal membranes in *C. elegans* (**Fig. 2F**). In wild-type animals, a short (10 min) incubation with fluorescein did not result in detectable fluorescence signals in the intestine (**Fig. 2G**). By contrast, the same procedure led to a dramatic accumulation of fluorescein in the intestine of *tsp-1* deficient animals (**Fig. 2G-2J**). This difference was notable at the L4 stage and particularly prominent in adults at day 5 post L4 (**Fig. 2J**). We observed similar defects in membrane barrier functions using RNA interference (RNAi) against two different coding regions of *tsp-1* (**fig. S5**). These results indicate that TSP-1 up-regulation during aging or by thermal stress may physiologically maintain intestinal barrier functions.

**Fig. 2.**
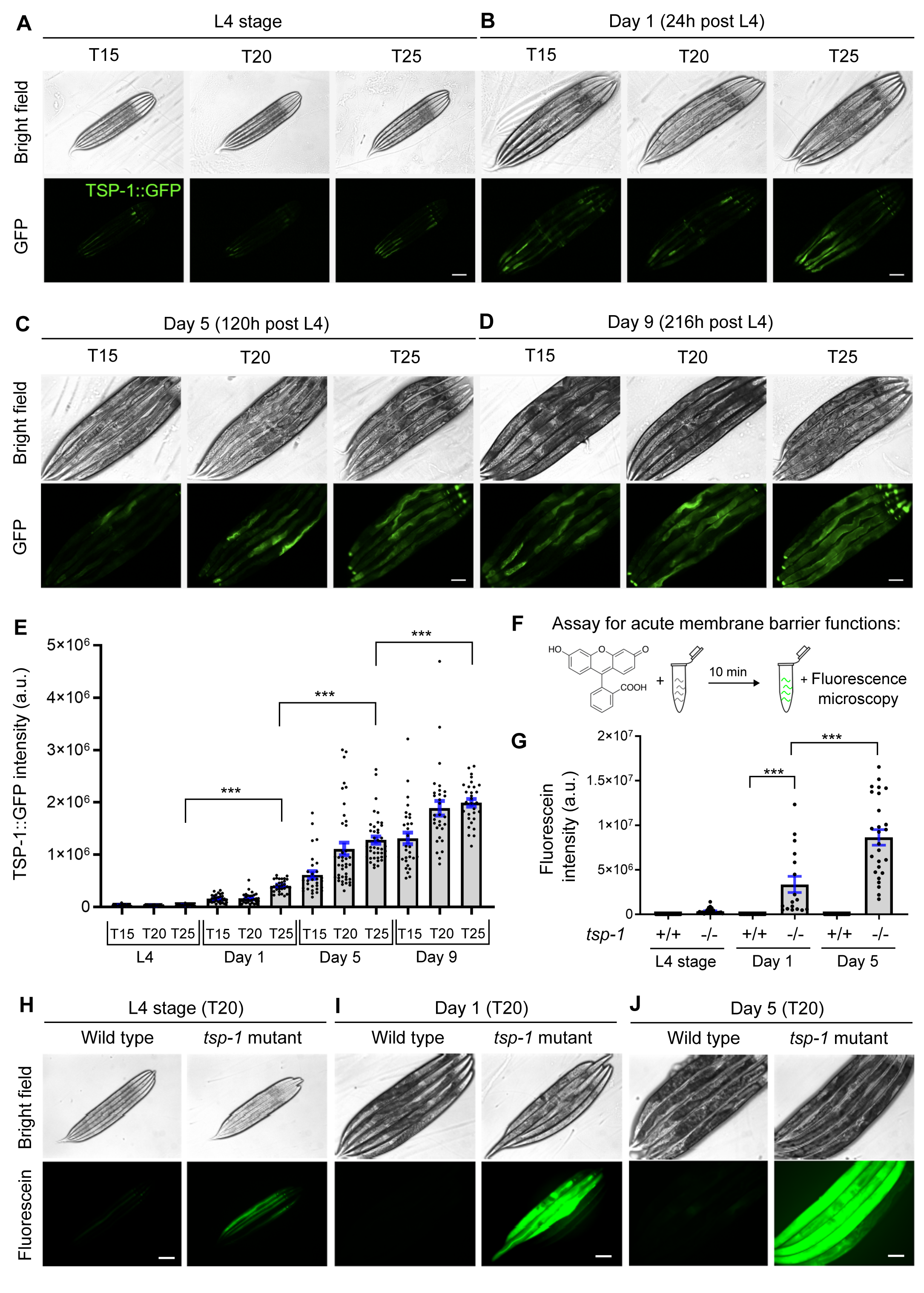
Expression and roles of TSP-1 in membrane barrier integrity in adults and during aging. (A-D) Representative epifluorescence images showing TSP-1::GFP up-regulation from L4 (A) to young adults (B) and during aging (C-D). (E) Quantification of fluorescence intensities of TSP-1::GFP under conditions indicated. *** indicates *P* < 0.001 (n > 30 animals per condition). (F) Schematic of the assay for acute membrane barrier functions. (G) Quantification of intensities of fluorescein accumulated by live animals under conditions indicated. *** indicates *P* < 0.001 (n > 20 animals per condition). (H-J) Representative epifluorescence images showing fluorescein accumulation in animals of indicated stages, temperature and genotypes (wild type and *tsp-1* deletion mutant allele *ok3594*). Scale bars: 100 µm.

We characterized how early-life thermal stress (ELTS, 28 °C for 24 hrs starting from L4 stages) may trigger lasting TSP-1::GFP expression. In control animals at 20 °C, TSP-1::GFP remained at low baseline levels largely unaltered for the first 48 hrs and became slightly elevated at 72 hrs (**Fig. 3A**). By contrast, ELTS (28 °C for 24 hrs at L4) triggered robust up-regulation of TSP-1::GFP abundance at 24 hrs post L4, which remained still high at 20 °C even for another 24 and 48 hrs after the initial ELTS (**Fig. 3A**). Chronic exposure to 28 °C starting at L4 induced stronger expression of TSP-1::GFP for 48 hrs than 24 hrs, but reached peak levels at 72 hrs (**Figs. 3B, C**). We took advantage of the robust TSP-1::GFP activation by ELTS and performed RNAi screens for genes that are required for TSP-1::GFP up-regulation. Given the importance of transcription factors and histone modifying enzymes in epigenetic gene regulation (*32–34*), we assembled a customized library of RNAi clones targeting >100 genes with adequate expression in the intestine (transcript per million >2) and that encode proteins including stress-responding transcription factors and chromatin/epigenetic regulators (**Fig. 3D, table S1**). From such candidate screens, using RNAi vector only as negative control and *tsp-1* RNAi as a positive control, we found that RNAi against only one hit, *cbp-1*, robustly prevented TSP-1::GFP up-regulation by ELTS (**Fig. 3E, 3F**). *cbp-1* encodes the *C. elegans* ortholog of histone acetyltransferase p300/CBP that promotes gene transcription (*35–43*). RNAi against many of the known genes encoding heat-responding transcription factor, including *hsf-1*, *hsf-2*, *hif-1* and *nhr-49*, did not block TSP-1::GFP up-regulation (**table S1**). Although TSP-1::GFP remained high for 24 and 48 hrs at 20 °C post initial ELTS (**Fig. 3C**), qRT-PCR measurements revealed that the up-regulation of *tsp-1* mRNA transcripts by 28 °C was transient but not sustained, and required CBP-1 (**Fig. 3F**). These results indicate that transient ELTS can trigger lasting up-regulation of TSP-1 abundance, which requires initial CBP-1-dependent *tsp-1* transcription but remains stable even after *tsp-1* expression has normalized after ELTS.

**Fig. 3.**
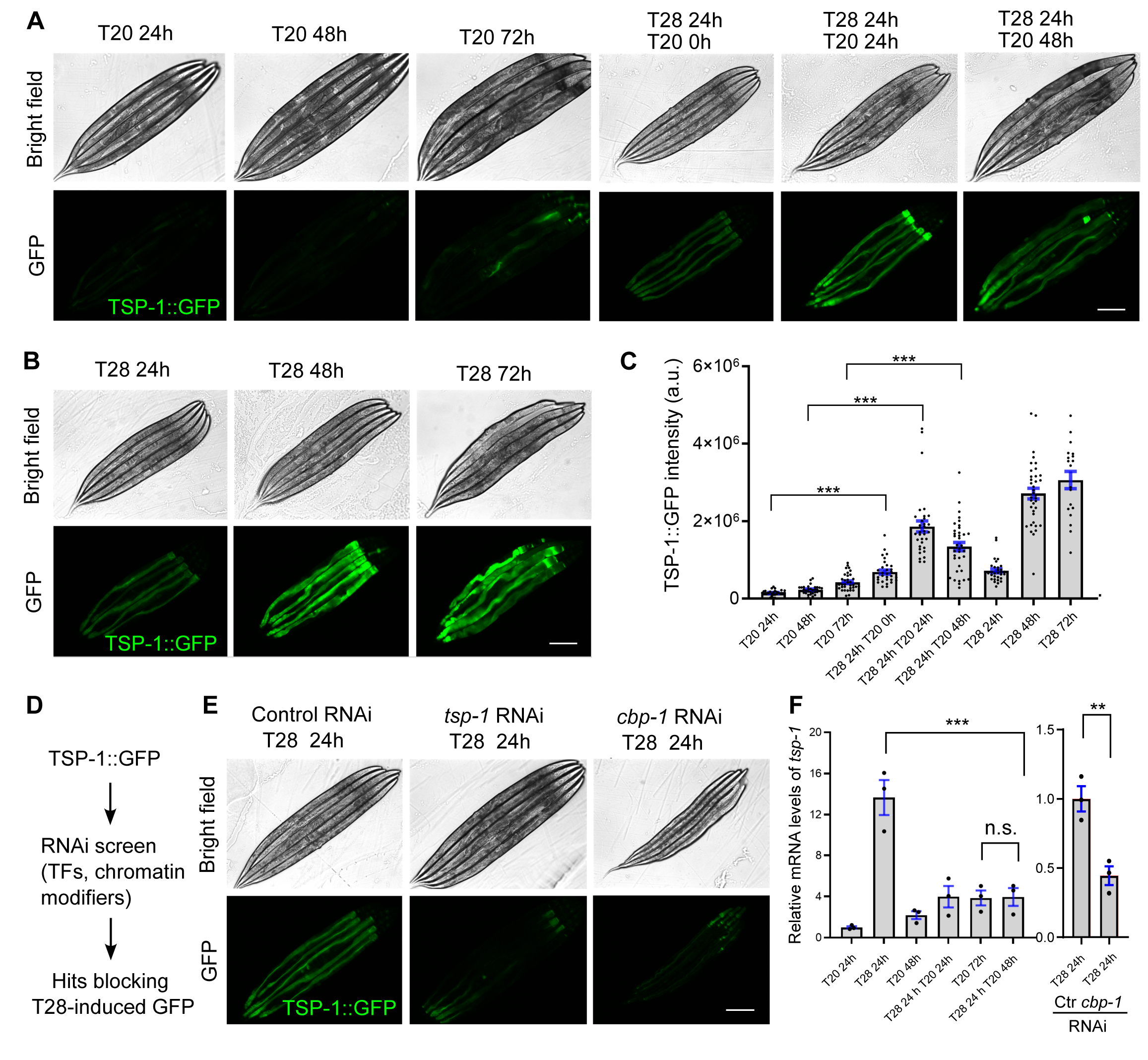
Early-life thermal stress-triggered TSP-1::GFP is long-lasting and requires CBP-1. (A) Representative epifluorescence images showing long-lasting up-regulation of TSP-1::GFP upon transient early-life thermal stress (ELTS, T28 for 24 hrs at L4). (B) Representative epifluorescence images showing up-regulation of TSP-1::GFP upon sustained thermal stress (T28 for 24, 48 or 72 hrs starting at L4). (C) Quantification of fluorescence intensities of TSP- 1::GFP under conditions indicated. *** indicates *P* < 0.001 (n > 30 animals per condition). (D) Schematic of experimental flow using RNAi screen to identify genes required for ELTS induction of TSP-1::GFP. (E) Representative bright-field and epifluorescence images showing that RNAi against *tsp-1* or *cbp-1* blocks up-regulation of TSP-1::GFP by ELTS. (F) Quantitative RT-PCR measurements of *tsp-1* expression levels under conditions indicated, showing transient *tsp-1* up-regulation by ELTS in a CBP-1-dependent manner. *** indicates *P* < 0.001 (three independent biological replicates). Scale bars: 100 µm.

We aimed to determine the subcellular localization and stability of the heat-induced TSP- 1 tetraspanin web formation. We generated strains with CRISPR-mediated wrmScarlet knock-in at the endogenous *tsp-1* locus that was genetically crossed with previously validated GFP reporters for intestinal subcellular structures, including Akt-PH::GFP that binds to PIP2/PIP3 at the inner leaflet of plasma membranes, or GFP::C34B2.10(SP12) that labels the endoplasmic reticulum membrane (ERm) (*21, 44, 45*). Confocal microscopic imaging revealed the proximity of Akt-PH::GFP and TSP-1::wrmScarlet throughout intestinal apical membranes (**Fig. 4A**), corroborated by quantitative fluorescent intensity correlation analysis (**Fig. 4B**). By contrast, ERm::GFP and TSP-1::wrmScarlet did not exhibit apparent co-localization (**Fig. 4C, 4D**). We further assessed the temporal dynamics of tetraspanin webs by imaging endogenous TSP- 1::wrmScarlet (**Fig. 4E**) and found that the tetraspanin web structure exhibited markedly consistent stability across the entire field and time scale of imaging (**Fig. 4F, Movie S1**). These results indicate that TSP-1-dependent tetraspanin webs, once induced by heat, form rather stable structures closely associated with the intestinal apical membrane.

**Fig. 4.**
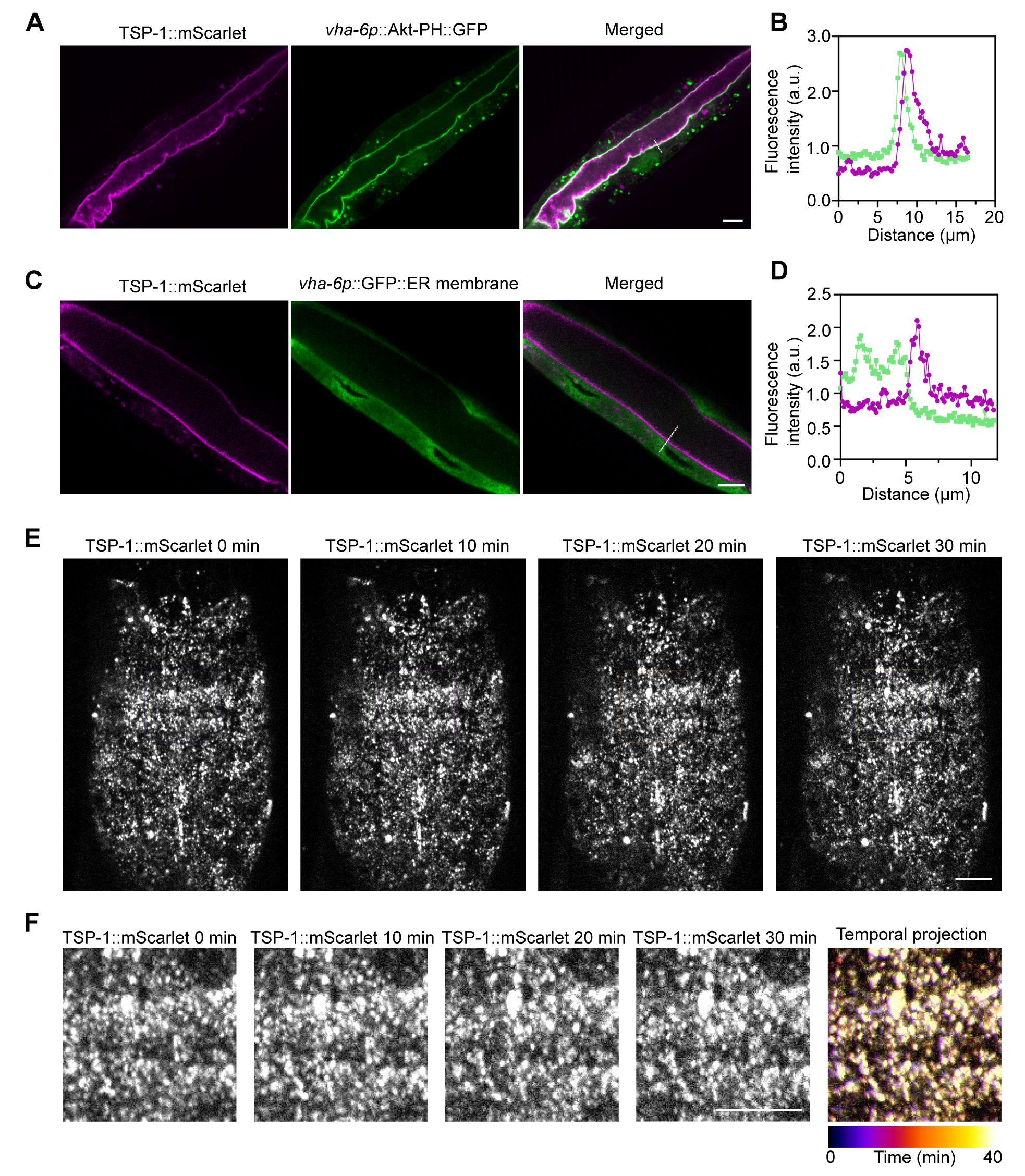
Subcellular localization and stability of tetraspanin web formed by endogenous TSP-1::wrmScarlet. (A) Representative confocal images showing co-localization of the marker Akt-PH::GFP, which binds to intestinal apical plasma membrane inner leaflet PIP2/3, with endogenous TSP-1 tagged with wrmScarlet by CRISPR. (B) Fluorescent intensity correlation analysis showing close proximity of Akt-PH::GFP and TSP-1::wrmScarlet. (C) Representative confocal images showing non-co-localization of the marker ERm::GFP, which labels intestinal ER membrane, with endogenous TSP-1 tagged with wrmScarlet by CRISPR. Scale bars: 10 µm. (D) Fluorescent intensity correlation analysis for ERm::GFP and TSP-1::wrmScarlet. (E) Representative confocal time-series images showing stability of tetraspanin webs formed by endogenous TSP-1::wrmScarlet, with enlarged views in (F). Scale bars: 10 µm.

Next, we investigated the consequences of TSP-1 up-regulation triggered by ELTS at the organismal level. To address this, we measured the population lifespan and survival rates of wild type and *tsp-1* deletion mutants under various heat stress conditions. Under constant 28 °C starting from L4, deficiency of *tsp-1* caused shortened lifespan and premature population death compared with wild-type animals (**Fig. 5A**). Transient ELTS (28 °C for 24 hrs at L4) extended the longevity of wild-type animals at subsequent normal cultivation temperature (20 °C). By contrast, the ELTS effect on longevity extension was abolished in *tsp-1* mutants (**Fig. 5B**). We applied 28 °C exposure to animals growing constantly for 48 hrs at 20 °C post L4, and found that transient ELTS (28 °C for 24 hrs at L4) enhanced survival rates in wild type but not *tsp-1*- deficient animals (**Fig. 5C**). We used RNAi against *cbp-1* (null mutations are lethal) and observed that reducing *cbp-1* expression caused even more shortened population lifespans (**Fig. 5D**). Given the more severe phenotype of *cbp-1* deficiency, CBP-1 likely regulates additional genes beyond *tsp-1*. Indeed, we found that the heat shock chaperon-encoding gene *hsp-16* was transiently induced by ELTS, and its induction required both HSF-1 and CBP-1 (**fig. S6**). Unlike *hsp-16*, ELTS induction of *tsp-1* is HSF-1 independent (**fig. S2**), yet its similarly transient induction leads to stable tetraspanin multimerization and web-like structural formation (**Fig. 3**). Finally, we used a CRISPR-mediated GFP knock-in allele of *cbp-1* to show that ELTS markedly increased the probability of nuclear entry by endogenous CBP-1 in the intestine, suggesting that CBP-1 plays an instructive (actively mediating the effect of heat) rather than permissive (gate-keeping to allow heat effect) role in *tsp-1* up-regulation by heat (**Fig. 5E, F**). Like induction of *tsp-1* mRNAs but not TSP-1 proteins, increased nuclear entry of CBP-1 also followed a similarly transient pattern upon ELTS (**fig. S7, Fig. 3**). Taken together, these results demonstrate that TSP-1 promotes survival under 28 °C heat stress and its CBP-1-dependent induction mediates effects of transient ELTS on lasting benefits to longevity and organismal resilience to subsequent heat stress conditions (**Fig. 5G**).

**Fig. 5.**
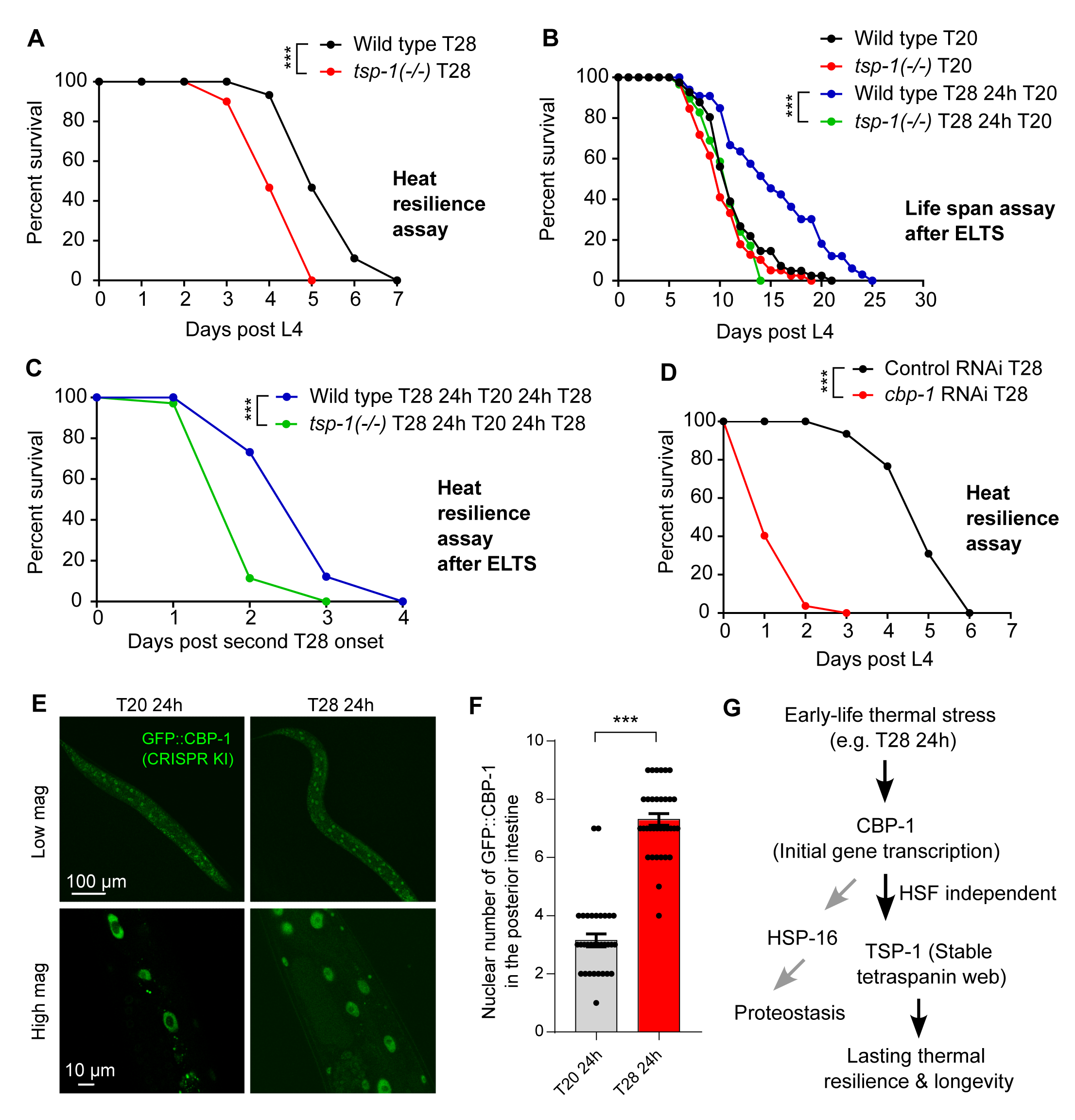
Early-life thermal stress-triggered thermal resilience and longevity requires TSP-1. (A) Lifespan curves of wild type and *tsp-1* mutants at 28 °C starting at L4.20% median and 28% maximal survival decrease in *tsp-1* mutants *** indicates *P* < 0.001 (n > 40 animals per condition). (B) Lifespan curves of wild type and *tsp-1* mutants at constant 20 °C or after transient 28 °C for 24 hrs at L4 (ELTS). 36% median and 20% maximal lifespan extension by ELTS in wild type, p=0.0008. *** indicates *P* < 0.001 (n > 35 animals per condition). (C) Lifespan curves of wild type and *tsp-1* mutants at 28 °C after transient 28 °C for 24 hrs at L4 (ELTS) . 33% median and 33% maximal survival decrease in *tsp-1* mutants *** indicates *P* < 0.001 (n > 35 animals per condition). (D) Lifespan curves of wild type animals with RNAi against *cbp-1* or control at 28 °C starting at L4. 80% median and 50% maximal survival decrease with RNAi against *cbp-1*. *** indicates *P* < 0.001 (n > 40 animals per condition) (E) Representative confocal fluorescence images showing ELTS-induced increase in numbers of nuclei with GFP::CBP-1 in the posterior intestine. (F) Quantification of numbers of nuclei with GFP::CBP-1 in the posterior intestine under conditions indicated. (G) Model of how ELTS induces lasting organismal thermal resilience through HSF-independent epigenetic regulation of TSP-1 and stable tetraspanin web structure formation.

To assess the sufficiency of TSP-1 in conferring stress resilience, we determined the gain-of-function effects of TSP-1 in *C. elegans* and ectopically in human cells. In *C. elegans*, we generated two transgenic strains carrying genomic *tsp-1* regions as extrachromosomal arrays at either low or high copy numbers (over-expression or OE, line 1 and 2, respectively) (**Fig. 6A**). qRT-PCR confirmed that *tsp-1* mRNA expression levels are differentially up-regulated in the two strains with array-positive animals (**Fig. 6A**). In thermal resilience assays, we found that OE line 2 with high-copy *tsp-1* arrays exhibited markedly enhanced survival rates upon continuous 28 °C heat stress, compared with array-negative control or OE line 1 with low-copy *tsp-1* arrays (**Fig. 6B**). In lifespan assays, OE line 2 exhibited similarly extended survival and longevity under 20 °C cultivation conditions, with a marked increase in both median (63%) and maximal (24%) lifespans in animals carrying high-copy expression of *tsp-1* (**Fig. 6C**). Furthermore, to assess the gain-of-function effect of TSP-1 in a heterologous cellular context, we over-expressed *C. elegans tsp-1* cDNA with a tagged *GFP* driven by the CMV promoter in human embryonic kidney cell lines (HEK293) (**Fig. 6D**). Such ectopic expression of *tsp-1* resulted in prominent tetraspanin web-like structures (**Fig. 6E**) and functionally protected HEK293 cells against thermal stress conditions (42 °C). These results demonstrate striking gain-of-function effects of tetraspanin-encoding *tsp-1* in both *C. elegans* and heterologous mammalian cells.

**Fig. 6.**
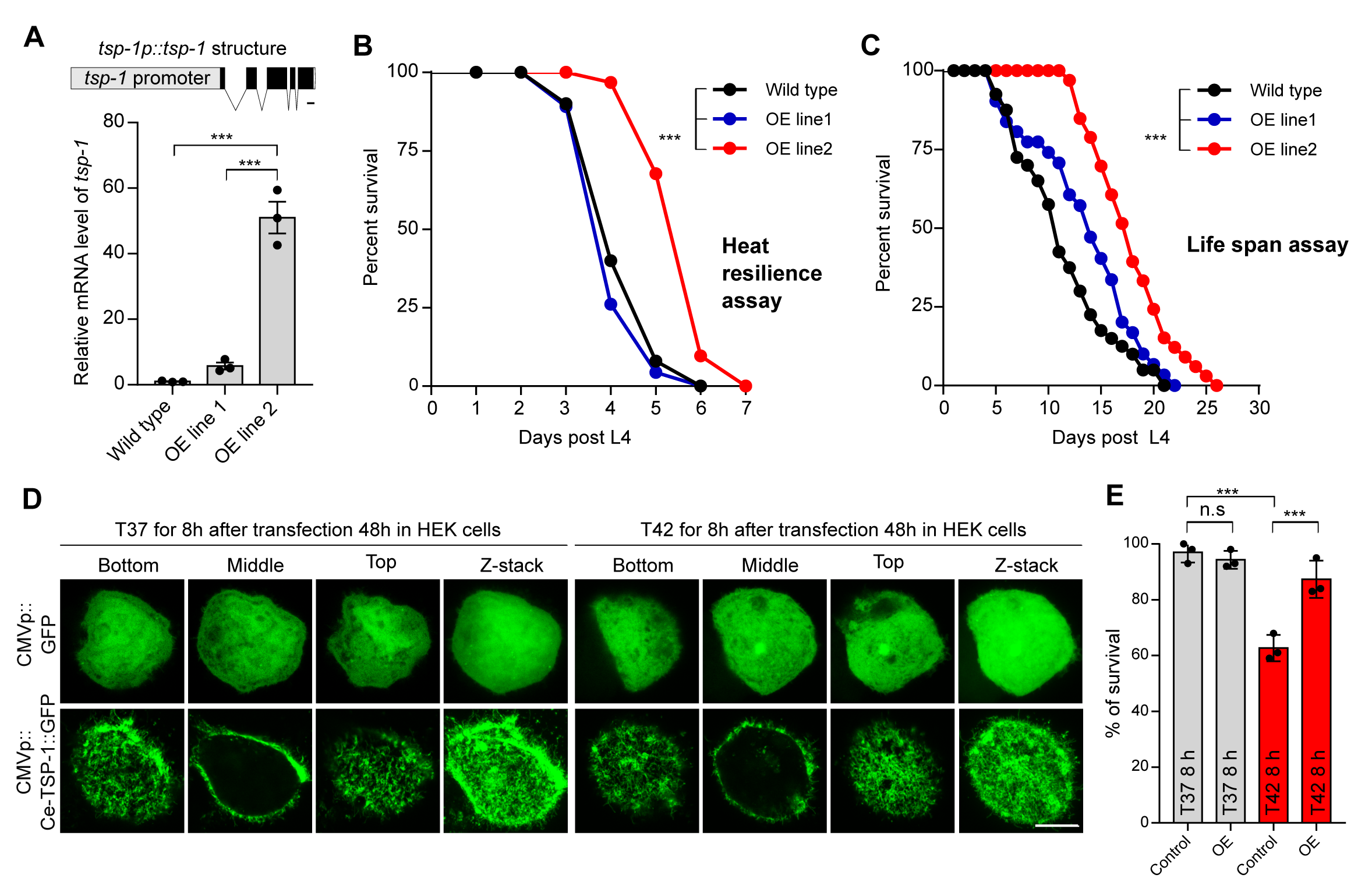
Gain-of-function of TSP-1 confers longevity extension in *C. elegans* and thermal resilience in human cells. (A) Schematic of the transgene that produces *tsp-1* gain-of-function over-expression lines (OE line 1 at low level, OE line 2 at high level as measured by qRT-PCR). *** indicates P < 0.001.(B) Lifespan curves of wild type and two representative *tsp-1* OE lines at 28 °C starting at L4. 50% median and 16% maximal survival extension by overexpression of tsp- 1 in wild type, *** indicates P < 0.001 (n > 40 animals per condition). (C) Lifespan curves of wild type and *tsp-1* OE lines at constant 20 °C. 63% median and 24% maximal lifespan extension by overexpression of tsp-1 in wild type, *** indicates P < 0.001 (n > 40 animals per condition). (D) Serial confocal fluorescence images showing tetraspanin web-like structures formed in the human cell line HEK293 by CMV promoter-driven expression of *C. elegans tsp-1::GFP* (bottom) but not *GFP* alone (top). Scale bars: 10 µm. (E) Quantification of survival rates after heat shock in HEK293 cells showing enhanced thermal resilience conferred by ectopic gain-of-function of *C. elegans* TSP-1::GFP. *** indicates *P* < 0.001 (three independent biological replicates).

## Discussion

Previous studies have shown that mild and transient metabolic or environmental stress in early life can produce beneficial effects, extending longevity in *C. elegans* (*38, 46–51*). While heat exposure activates numerous genes involved in proteostasis and defense responses, specific HSF-independent target genes with causal effects on longevity remain unidentified. In this work, we identify *tsp-1* as a heat-inducible gene that mediates effects of ELTS on organismal resilience and longevity. Regulatory mechanisms governing how heat stress activates CBP-1 in coordination with other unidentified factors await further exploration. Our data suggest that *tsp-1* expression leads to TSP-1 protein multimerization and the formation of stable tetraspanin web structures, which persist even in the absence of initial stimuli and *tsp-1* transcript up-regulation. This tetraspanin web-based stable protein structure formation represents a novel mechanism of cellular memory, distinct from previously known modes of epigenetic regulation primarily occurring in the nucleus, such as DNA and histone modifications.

We demonstrate that TSP-1 is crucial for maintaining intestinal membrane barrier functions and enhancing animal adaptation and survival under heat-stress conditions and during aging. The *C. elegans* genome encodes 21 tetraspanin family proteins, including *tsp-1* and *tsp-2* located in the same operon, which may be similarly regulated by heat. Other tetraspanin family members have been implicated in plasma membrane repair, extracellular vesicle formation, and oxidative stress responses (*52–55*). It is plausible that different tetraspanins may be regulated in a tissue-specific, developmental stage-specific, and stress-specific manner to confer distinct biological functions. In addition to heat that impacts membrane barrier and integrity, other types of specific membrane-perturbing stress can also induce *tsp-1* expression through CBP-1, which likely regulates additional transcriptional targets that together safeguard intestinal barrier functions (**fig. S8, 9**). Several mammalian tetraspanins have been shown to play important roles in maintaining blood-brain or retina-blood barriers in endothelial cells and attenuating inflammation, cell senescence, and organismal aging (*56–58*). Thus, we propose that the functional roles of tetraspanins are evolutionarily conserved, likely mediating long-lasting effects of transient cellular and organismal stresses in mammals, including humans.

## Materials & Methods

### C. elegans strains

1. *C. elegans* strains were grown on nematode growth media (NGM) plates seeded with *Escherichia coli* OP50 at 20°C with laboratory standard procedures unless otherwise specified. The N2 Bristol strain was used as the reference wild type (*59*). Mutants and integrated transgenes were back- crossed at least 5 times. Genotypes of strains used are as follows: *dmaIs125 [tsp-1p::tsp-1::GFP; unc-54p::mCherry]*, *dmaIs8 [hsp-16p::GFP; unc-54p::mCherry]; him-5(e1490)*, *hjIs14 [vha- 6p::GFP::C34B2.10(SP12 ER membrane) + unc-119(+)]*, *etEx2 [glo-1p::GFP::ras-2 CAAX* + *rol- 6], tsp-1(ok3594*), *hsf-1(sy441)*, *tsp-1(syb7456[tsp-1::wrmScarlet]*), *cbp-1(ust475[GFP::cbp-1])*.

For CRISPR knock-in of *cbp-1* with *3xFLAG::GFP::cbp-1*, a *cbp-1* promoter region was PCR-amplified with the primers 5′- gggtaacgccagCACGTGtgggctgactcgtgctg -3′ and 5′- ccgtcatggtctttgtagtctggtggttcatccatcaattagta -3′ from N2 genomic DNA. A *cbp-1* coding sequence region was PCR amplified with the primers 5′- AAGGAGGTGGAGGTGGAGCTatggatgaaccaccatcaa -3′ and 5′- cagcggataacaatttcacaatgcatcattggatatccacc -3′ from N2 genomic DNA. A 3xFLAG::GFP region was PCR-amplified with the primers 5′- gactacaaagaccatgacgg -3′ and 5′- AGCTCCACCTCCACC -3′ from SHG1890 genomic DNA. ClonExpress MultiS One Step Cloning Kit (Vazyme C113-02, Nanjing) was used to connect these fragments with vector amplified with 5′- tgtgaaattgttatccgctgg -3′ and 5′- caCACGTGctggcgttacc -3′ from L4440. The injection mix contained pDD162 (50 ng/ul), *cbp-1* repair plasmid (50 ng/ul), pCFJ90 (5 ng/ul), and three sgRNAs (30 ng/ul). The mix was injected into young adult N2 animals, and the coding sequence of 3xFLAG::GFP was inserted after the start codon using the CRISPR/Cas9 system. The *tsp-1::wrmScarlet* knock-in allele was generated similarly (Sunybiotech), with *wrmScarlet* sequences inserted before the termination codon of *tsp-1* in the N2 background.

For overexpression of *tsp-1* (3.26 kb), the *tsp-1* promotor (1.98 kb) and 1,283 bp genomic DNA fragment of *tsp-1* coding region was PCR-amplified with the primers 5′- aaatataatttcagggatgtggtctcaaat -3′ and 5′- tgacaagggtactgtagttcgtct -3′ from N2 genomic DNA. The injection mix contained *tsp-1p::tsp-1* (10-50 ng/µl) and *unc-54*p::GFP (25 ng/µl) to establish transgenic strains carrying extrachromosomal arrays of varying copy numbers.

### Compound and confocal imaging

Epifluorescence compound microscopes (Leica DM5000 B Automated Upright Microscope System) were used to capture fluorescence images (with a 10× objective lens). Animals of different genotypes and different stages (L4, Day1, Day5, Day9) and different heat treatment were randomly picked and treated with 10 mM sodium azide solution (71290-100MG, Sigma-Aldrich) in M9, aligned on an 2% agarose pad on slides for imaging. The same settings (for bright-field: exposure time 1 second, for GFP: exposure time 10 seconds) were maintained for the images of all samples. The integrated density (IntDen) of TSP-1::GFP was measured by NIH image program (Fiji image J) , average of mean gray value (three background area of each image randomly selected) were employed to normalized the TSP-1::GFP. For confocal images, the worms were randomly chosen and treated with 10 mM sodium azide in M9 solution and aligned on an 2% agarose pad on slides and images acquired using a confocal microscope (Leica TCS SPE) with a 63× objective lens, the same settings were maintained for the images of all samples. Imaging in Figure 4 was conducted under room temperature conditions using a Nikon Eclipse Ti inverted microscope equipped with a Borealis beam conditioning unit (Andor), a CSU-W1 Yokogawa spinning disk (Andor; Belfast, Northern Ireland), a 100X PlanApo TIRF 1.49 numerical aperture (NA) objective (Nikon; Toyko, Japan), an iXon Ultra EMCCD camera (Andor), and a laser merge module (LMM5, Spectral Applied Research; Exton, PA) containing 405, 440, 488, and 561-nm laser lines. Micro-Manager (UCSF) was used to control all the hardware. Fiji (NIH) (*60*) and Prism (Graphpad software, Inc) were used for image display and co-localization analysis.

### SDS-page and native-page western blotting

For SDS-PAGE samples, stage-synchronized animals for control and experiment groups were picked (n = 50) in 60 μl M9 buffer and lysed directly by adding 20 μl of 4x Laemmli sample buffer (1610747, Bio-Rad) contain 10% of 2-Mercaptoethanol (M6250-100ML, Sigma(v/v)). Protein extracts were denatured at 95 °C for 10 min and separated on 10% SDS-PAGE gels (1610156, Bio-Rad) at 80 V for ∼40 min followed by 110 V for ∼70 min. The proteins were transferred to a nitrocellulose membrane (1620094, Bio-Rad,) at 25 V for 40 mins by Trans-Blot® Turbo™ Transfer System (Bio-Rad). The NC membrane was initially blocked with 5% nonfat milk and 2% BSA (A4503, Sigma (v/v)) in tris buffered saline with 0.1% Tween 20 (93773, Sigma) (TBS-T) at room temperature for 1 h. Proteins of interest were detected using antibodies against GFP (A6455, Invitrogen) and Histone H3 (ab1791, Abcam) in cold room for overnight. After three washes of 5 min each with tris-buffered saline with 0.1% Tween 20, anti-rabbit IgG, HRP-linked Antibody (7074S, Cell Signaling Technology) was added at a dilution of 1:5000.

For native-PAGE samples, stage-synchronized animals were washed down from NGM plates using M9 solution and subjected to 850 *g* for 60 seconds, and the pellet animals were resuspended in pre-cooled 300 µl lysis buffer (M9 buffer + Protein inhibitors cocktails, A32965, Thermo Fisher), then lysed by TissueRuptor (Motor unit ‘6’) for 10 seconds and taken out, repeated 3-5 times, followed by diluting the samples with Native Sample Buffer (161-0738, Bio-Rad). Proteins were resolved by 4–15% Mini-PROTEAN® TGX™ Precast Protein Gels (4561086, Bio-Rad,) and transferred to a nitrocellulose membrane (1620094, Bio-Rad). The NC membrane was initially blocked with 5% nonfat milk and 2% BSA (A4503, Sigma (v/v)) in Tris buffered saline with 0.1% Tween 20 (93773, Sigma) (TBS-T) at room temperature for 1 h. native TSP-1::GFP were detected using antibodies against GFP (1:1000, 66002-1-Ig, Proteintech) at cold room overnight. After three washes of 5 min each with Tris-buffered saline with 0.1% Tween 20, goat anti-Mouse IgG (H+L) Secondary Antibody [HRP] (NB7539, Novus) added at a dilution of 1:5000 as the secondary antibody.

### Fluorescein assay

Stage-synchronized animals (L4, Day1 and Day5) of different genotypes at 20°C were randomly picked to 20 μl M9 solution in 1.5 mL tube, followed by treatment with 100 μl of Fluorescein (F0095-25G, TCI America, 10 mg/ml in M9 buffer) at room temperature for 10 minutes. Worms were transferred to a NGM plate (without OP50), collected to new NGM plate with 200 μl M9 buffer to wash for three times, followed by randomly picking and treatment with 10 mM sodium azide in M9 buffer and aligned on an 2% agarose pad on slides for compound microscope imaging. The integrated density (IntDen) of fluorescein was measured by NIH image program (Fiji image J), average of mean gray value (three background areas of each image randomly selected) was employed to normalize the fluorescein signals.

### RNA interference (RNAi) cloning and screen

To clone the *tsp-1* RNAi sequences, *tsp-1* exon 1-3 (Chr. III, C02F5.8, 1124 bp size, primer forward:

GGCGGCCGCTCTAGAACTAGTACAGATTTCCTCCACCTCTTCC, primer reverse: TCCACGCGTCACGTGGCTAGCATTCCTAATTTTTCAGAGCCCACC) and exon 1-5 ( Chr. III, C02F5.8, 1472 bp size, primer forward: GGCGGCCGCTCTAGAACTAGTTCCACCTCTTCCACCTTCATTAC, primer reverse: TCCACGCGTCACGTGGCTAGCTGACAAGGGTACTGTAGTTCGTCT) were PCR-amplified from wild-type N2 gDNA and subcloned into the SpeI and NheI sites of a pL4440 expression vector with NEBuilder® HiFi DNA Assembly Cloning Kit (E2621L, NEB). Briefly, PCR was performed with the following protocol on a MyCyclerTM Thermal Cycler (Bio–Rad): 98 °C for 30 s, 98 °C for 10 s, 55 °C for 30 s, 72 °C for 2 min (35 cycles); 72 °C for 5 min, and a final hold at 12 °C. The PCR products were analyzed on a 1.5 % agarose gel. 25 ng of precut pL4440 vector with 2-fold excess of *tsp-1* PCR fragment (50 ng) were used for assembly in a thermocycler at 50°C for 30 minutes, 25 ng of precut pL4440 vector with same water instead of PCR fragment were employed as native assembly control. 5 μl of assembly preparations were transformed to NEB 5-alpha Competent *E. coli* (C2987H, NEB) by heat shock at exactly 42°C for exactly 30 seconds. Positive clones were verified by bacteria PCR by 2x thermo scientific dream taq green PCR master mix (pL4440 forward: CTTATCGAAATTAATACG, pL4440 reverse: AGGGCGAATTGGGTACCG). Positive pL4440-*tsp-1* RNAi plasmids were transformed to competent HT115 by electroporation and verified the positive HT115 clones by bacteria PCR.

RNAi and screen for hits blocking TSP-1::GFP were performed by feeding worms with *E. coli* strain HT115 (DE3) expressing double-strand RNA (dsRNA) targeting endogenous genes. Briefly, dsRNA–expressing bacteria were replicated from the Ahringer library to LB plates containing 100 μg/ml ampicillin (BP1760-25, Fisher Scientific) at 37 °C for 16 hrs. Single clone was picked to LB medium containing 100 μg/ml ampicillin at 37 °C for 16 hrs and positive clones (verified by bacteria PCR with pL4440 forward and pL4440 reverse primers) were spread onto NGM plates containing 100 μg/ml ampicillin and 1 mM isopropyl 1-thio-β-Dgalactopyranoside (IPTG, 420322, Millopore) for 24 hrs (namely RNAi plates). Developmentally synchronized embryos from bleaching of gravid adult hermaphrodites were plated on RNAi plates and grown at 20 °C to L4 stage followed by transfer to 28 °C incubator for 24 hrs. Randomly selected population was observed under the epifluorescence microscope (SMZ18, Nikon) with hits considered to block TSP-1::GFP when observing GFP levels comparable to that by *tsp-1* RNAi (positive control).

### qRT-PCR

Animals were washed down from NGM plates using M9 solution and subjected to RNA extraction using TissueDisruptor and RNA lysis buffer (Motor unit ‘6’ for 10 seconds and take it out, repeat 3-5 times on ice) and total RNA was extracted following the instructions of the Quick-RNA MiniPrep kit (Zymo Research, R1055) and reverse transcription was performed by SuperScript™ III (18080093, Thermo scientific). Real-time PCR was performed by using ChamQ Universal SYBR qPCR Master Mix (Q711-02, Vazyme) on the Roche LightCycler96 (Roche, 05815916001) system. Ct values of *tsp-1* were normalized to measurements of *rps-23* (*C. elegans*) levels. Primer for qRT-PCR: *tsp-1* (forward, CTTTGATTGCCGTTGGATTT; reverse, CCCAAAGAAAGGCCGATAAT), *hsp-16.2* (forward: ACGTTCCGTTTTTGGTGATCTTAT; reverse, TCTGGTTTAAACTGTGAGACGTTG) and *rps-23* (forward, CGCAAGCTCAAGACTCATCG; reverse, AAGAACGATTCCCTTGGCGT).

### Thermal resilience and lifespan assays

Animals were cultured under non-starved conditions for at least 2 generations before heat stress assays. For treatment of “early-life thermal stress” (ELTS), bleach-synchronized eggs were growth to the L4 stage at 20°C, and populations were kept at 28 °C for 24 hrs and then recovered for 0-72 hrs at 20 °C. For thermal resilience assays, stage-synchronized L4 stage worms (n > 30) were picked to new NGM plates seeded with OP50 and transferred to 28 °C incubator. Animals were scored for survival per 24 hrs. Worms failing to respond to repeated touch of a platinum wire were scored as dead. For lifespan assays, stage-synchronized L4 stage worms (n = 50) were picked to new NGM plates seeded with OP50 containing 50 μM 5-fluoro-2′-deoxyuridine (FUDR) to prevent embryo growth at 20 °C incubator. Animals were scored for survival per 24 hrs. Worms failing to respond to repeated touch of a platinum wire were scored as dead.

### Mammalian expression constructs for *C. elegans* TSP-1

The *C. elegans tsp-1* open reading frame was PCR-amplified with the primers 5′- GCGGCCTTAATTAAACCTCTAGAATGGCAACTTGGAAATTTATCATACG -3′ and 5′- AGCTCGAGATCTGAGTCCGGcAAAACGAGTGTCTTCGGTGATG -3′ from cDNA prepared from heat-induced (T28 24 hrs) wild-type animals. GFP fragment was PCR-amplified with 5′- gCCGGACTCAGATCTCGAGCTATGGTGAGCAAGGGCGCCG -3′ and 5′- CGGATCTTACTTACTTAGCGGCCGCTTACTTGTACAGCTCATCCATGCC -3′ from the pHAGE2-gfp plasmid. The *tsp-1*-GFP fragment was PCR-amplified using overlapping PCR and sub-cloned by T4 DNA Ligase to the pHAGE2-gfp plasmid at the XbaI and NotI sites.

### HEK293T cells and thermal resilience assay

Human embryonic kidney (HEK) 293T cells were maintained in Dulbecco’s modified Eagle’s medium with 10% inactive fetal bovine serum and penicillin-streptomycin (Gibco, Grand Island, 15140122) at 37 °C supplied with 5% CO2 in an incubator (Thermo Fisher Scientific) with a humidified atmosphere. Cells were washed once using PBS and digested with 0.05% trypsin-EDTA (Gibco) at 37 °C for routine passage of the cells. All HEK 293T cells were transiently transfected with indicated constructions using the lipo2000 (1 mg/ml, LIFE TECHNOLOGIES) reagent. The lipo2000/DNA mixture with the ratio of lipo2000 to DNA at 3:1 was incubated for 30 min at room temperature before being added to the HEK 293T cell cultures dropwise. For thermal resilience assay, mock control and transfected HEK293T cells (48 h) in 24 well plate were placed in a culture incubator with an ambient temperature at 42°C and humidified 5% CO2 for 8 h followed by cell death assay or imaging with 4% PFA treatment for 12 min at room temperature. For cell death assay, the collected cells were resuspended with 100 μl buffer with addition of 0.1 μl Sytox blue (Thermo Fisher Scientific) for an additional 15 min at room temperature. 25 ul of incubated cells were loaded into ArthurTM cell analysis slide (Nanoentek, AC0050). The fluorescence intensity was measured for individual cells using automated cytometry (ArthureTM image based cytometer, Nanoentek, AT1000) by viability assay. The 226 RFU (Fluorescence) threshold and cell size min 5 to max 25 were used for cell death analysis and quantification.

## Statistics

Data were analyzed using GraphPad Prism 9.2.0 Software (Graphpad, San Diego, CA) 389 and presented as means ± S.D. unless otherwise specified, with P values calculated by unpaired two-tailed t-tests (comparisons between two groups), one-way ANOVA (comparisons across more than two groups) and two-way ANOVA (interaction between genotype and treatment), with post-hoc Tukey HSD and Bonferroni’s corrections. The lifespan assay was quantified using Kaplan–Meier lifespan analysis, and *P* values were calculated using the log-rank test.

## Acknowledgments

Some strains were provided by *the Caenorhabditis* Genetics Center (CGC), which is funded by the NIH Office of Research Infrastructure Programs (P40 OD010440).

## Author Contributions

Conceptualization: WIJ, ODW, DKM

Methodology: WIJ, HDB, BW, AW, MK, FO, JD, XH, SG Investigation: WIJ, HDB, BW, AW, MK, FO, JD, XH, SG, ODW, DKM

Visualization: WIJ, HDB, ODW, DKM Supervision: ODW, DKM Writing—original draft: WIJ, DKM

Writing—review & editing: WIJ, HDB, BW, AW, MK, FO, JD, XH, SG, ODW, DKM

## Data and Materials Availability

All data needed to evaluate the conclusions in the paper are present in the paper and/or the Supplementary Materials.

---

**fig. S1.**
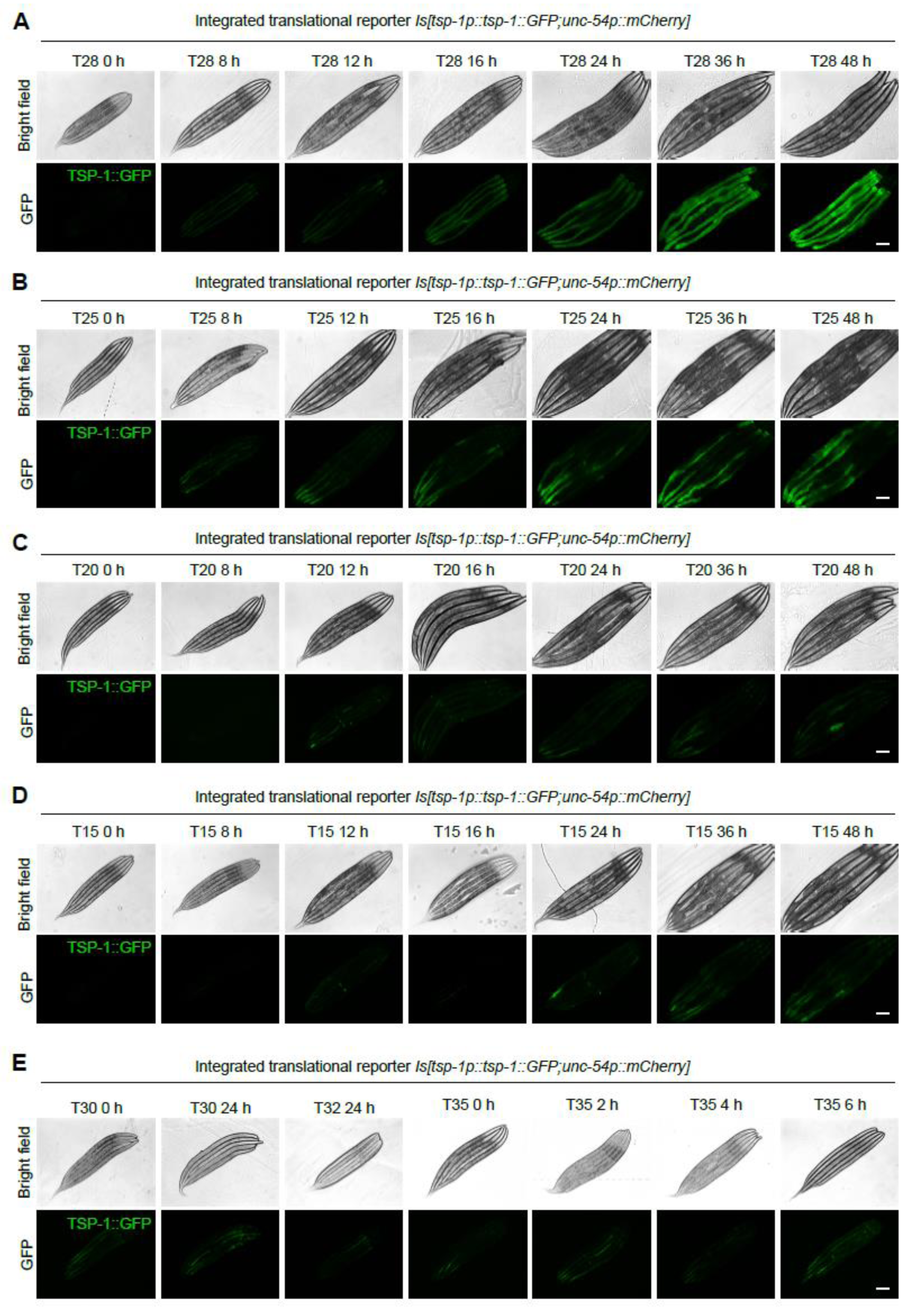
Duration and temperature-dependent induction of TSP-1::GFP. (A-E) Representative bright-field and epifluorescence images showing expression of *tsp-1p::tsp- 1::GFP* under temperatures and durations indicated. Scale bars: 100 µm.

**fig. S2.**
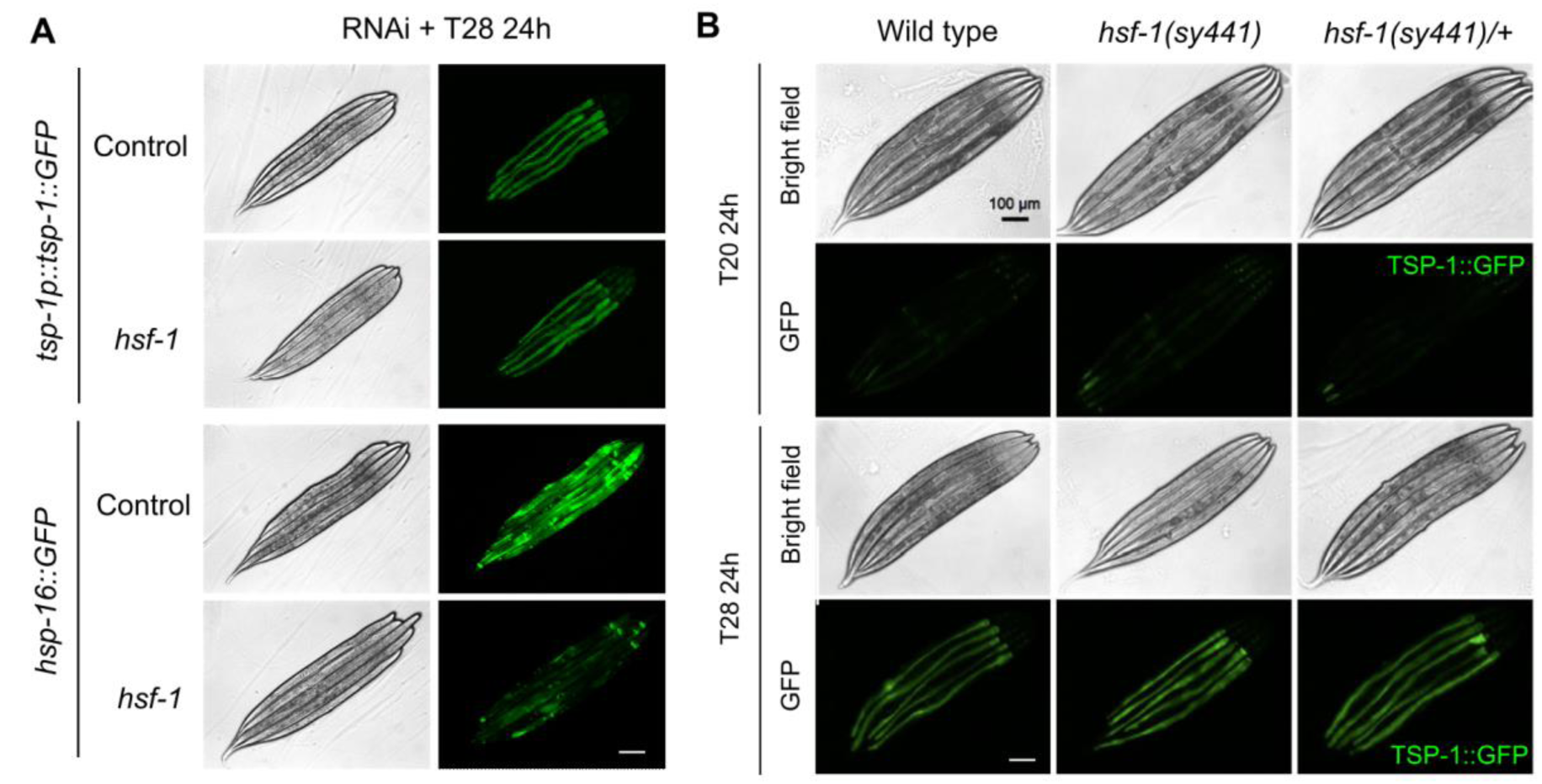
TSP-1 induction by T28 is independent of HSF-1. (A) Representative bright-field and epifluorescence images showing expression of *tsp-1p::tsp-1::GFP* or *hsp-16p::GFP* at 28 °C for 24 hrs, with control and RNAi against *hsf-1*. (B) Representative bright-field and epifluorescence images showing expression of *tsp-1p::tsp-1::GFP* at 28 °C for 24 hrs in wild type, *hsf-1(sy441)* reduction-of-function heterozygous or homozygous mutants. Scale bars: 100 µm.

**fig. S3.**
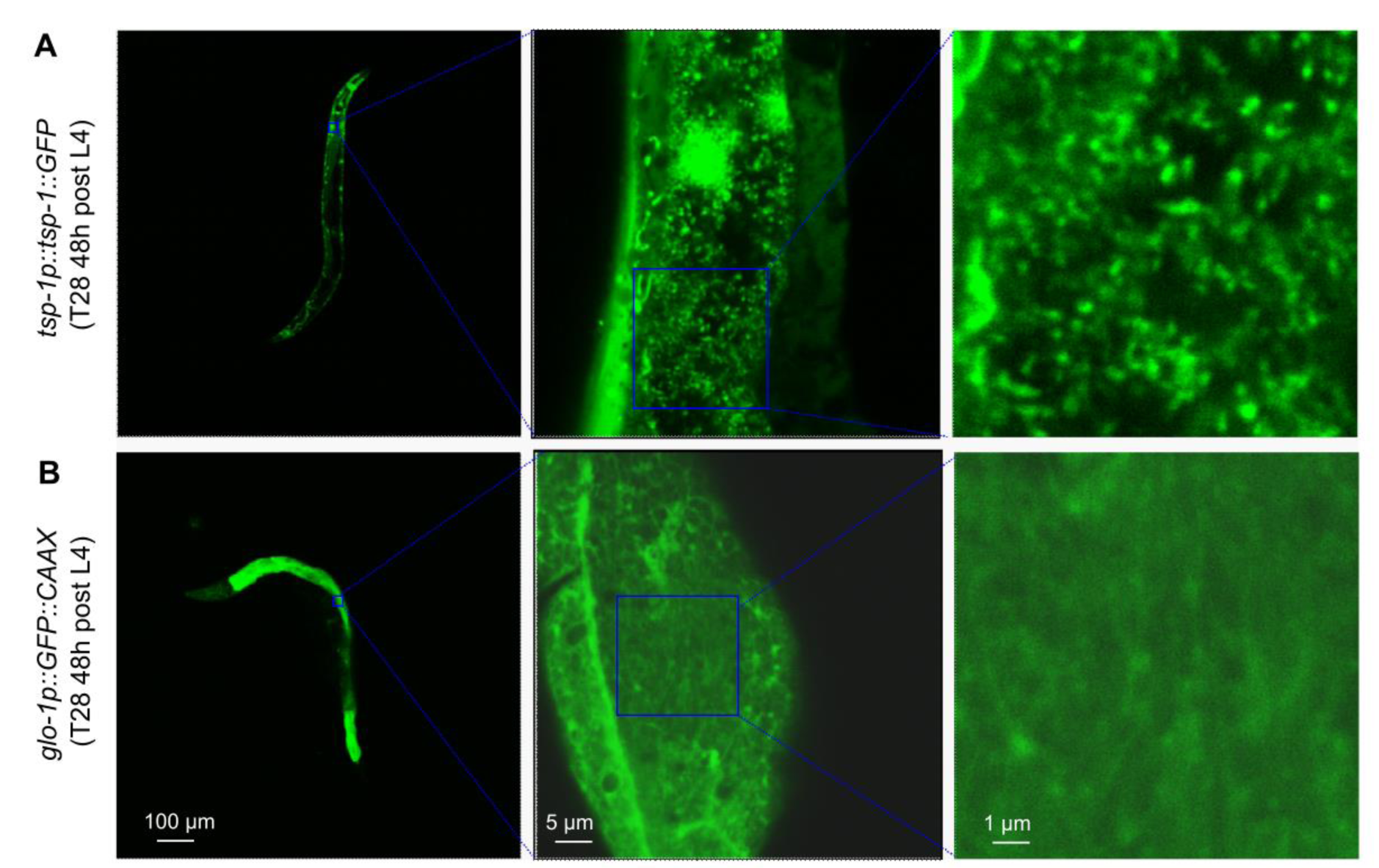
Tetraspanin web regulation by heat is specific for TSP-1. (A) Representative confocal fluorescence images showing T28-induced tetraspanin web structure formation by *tsp-1p::tsp- 1::GFP* transgenes. (B), Representative confocal fluorescence images showing intestinal membrane GFP from *glo-1p::GFP::CAAX* under identical conditions (T28 48 hrs).

**fig. S4.**
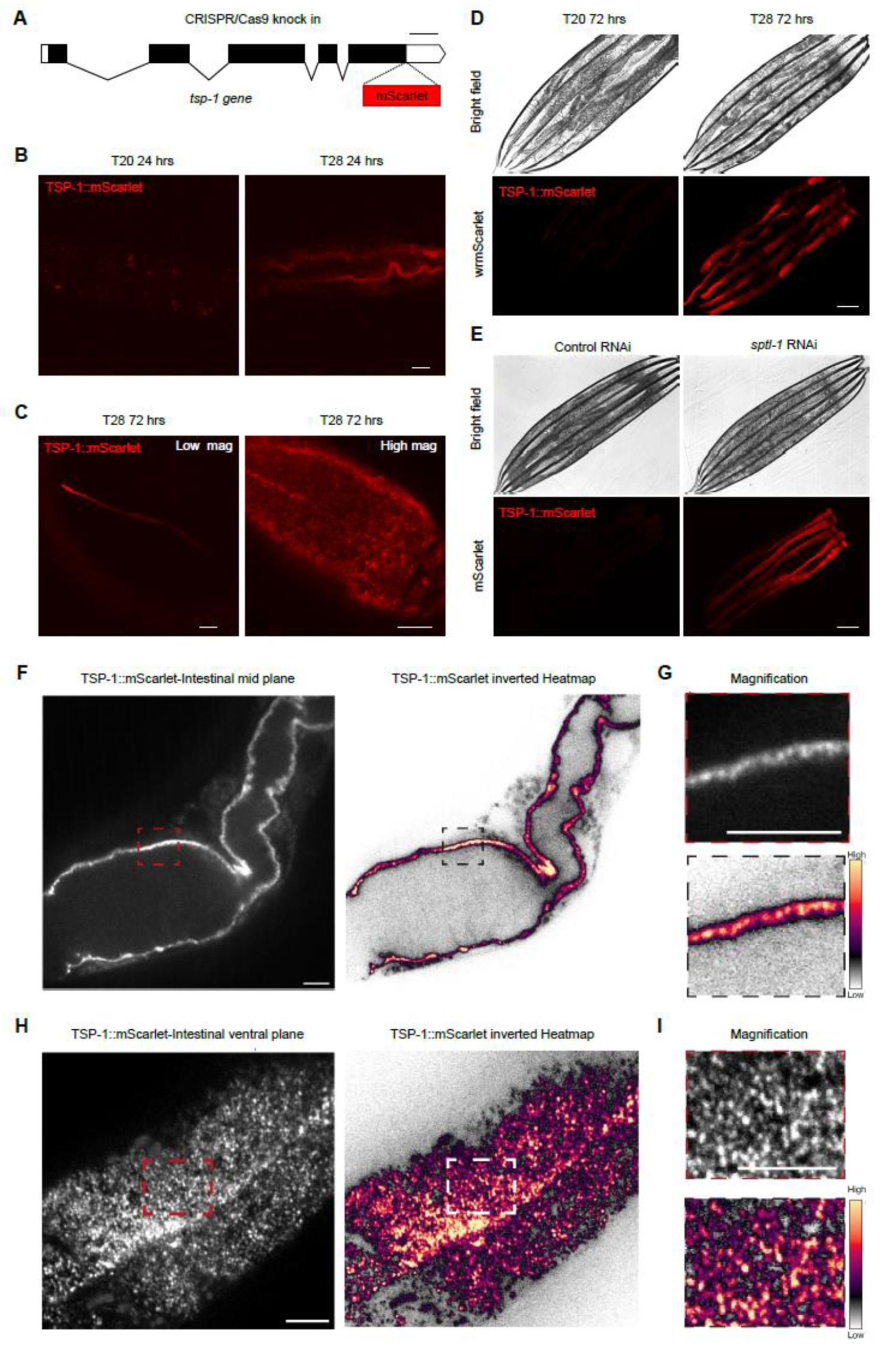
Heat-induced *tsp-1* endogenously tagged with wrmScarlet. (A) Schematic of tsp-1 gene structure showing CRISPR-mediated knock-in of wrmScarlet at the C-terminus of TSP-1. (B) Representative confocal fluorescence images showing up-regulation of endogenous TSP- 1::wrmScarlet by 28 °C for 24 hrs. (C) Representative confocal fluorescence images showing low and high-mag views of endogenous TSP-1::wrmScarlet induced by 28 °C for 72 hrs. (D)Representative epifluorescence images showing up-regulation of endogenous TSP- 1::wrmScarlet by 28 °C for 72 hrs. (E) Representative epifluorescence images showing up- regulation of endogenous TSP-1::wrmScarlet by RNAi against *sptl-1*, loss of which disrupts the biosynthesis of sphingolipids mimicking heat-induced membrane effects. (F-I) Representative spinning disc confocal images showing high-resolution views of tetraspanin web structures.

**fig. S5.**
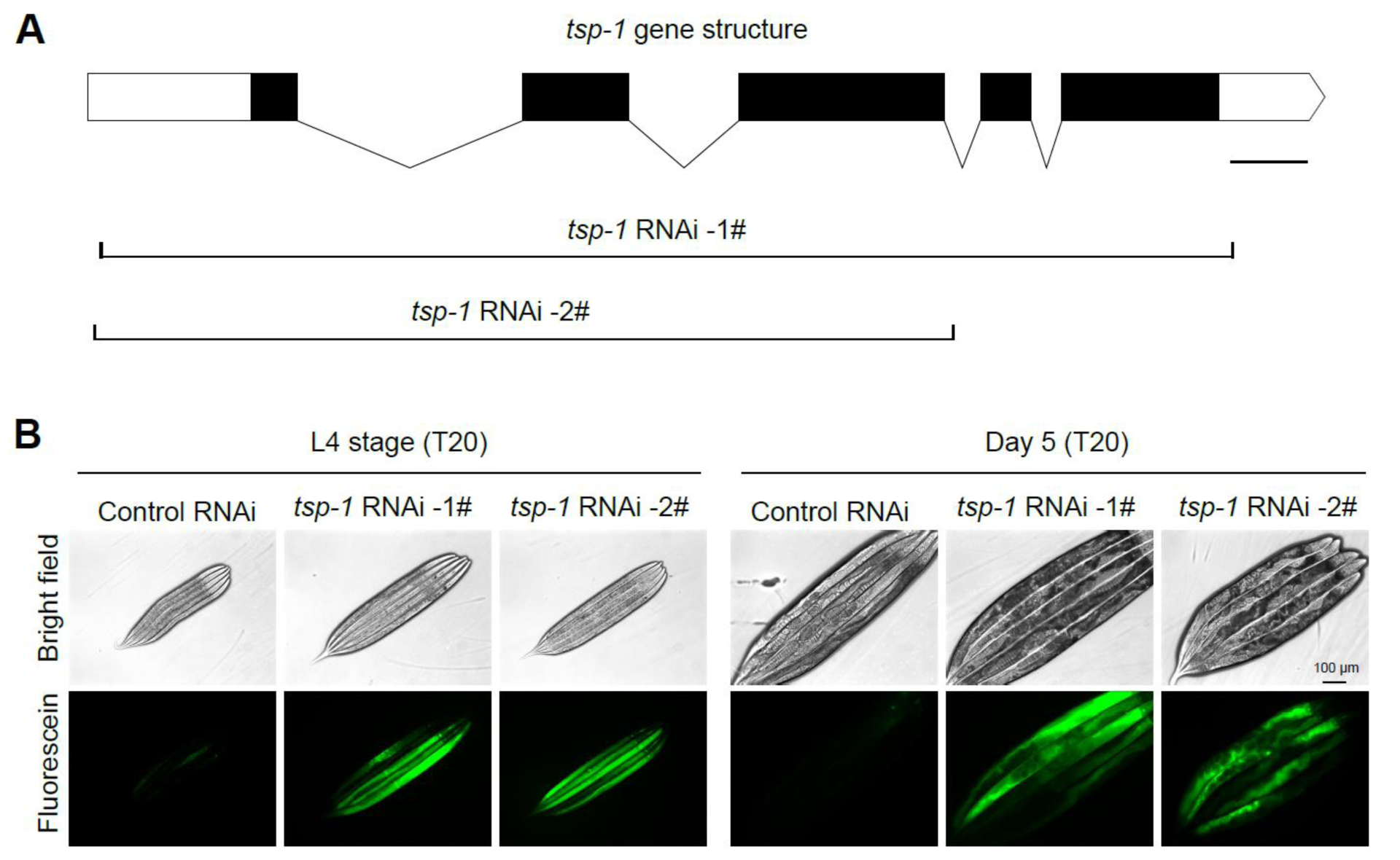
RNAi against *tsp-1* recapitulates mutant phenotype in membrane barrier functions. (A) Schematic of *tsp-1* gene structure showing two genomic regions used to construct RNAi for expression in *E. Coli* and feeding to *C. elegans*. (B) Representative epifluorescence images showing enhanced fluorescein uptake in *tsp-1* RNAi-treated animals, at both L4 stage and Day 5- old animals. Scale bars: 100 µm.

**fig. S6.**
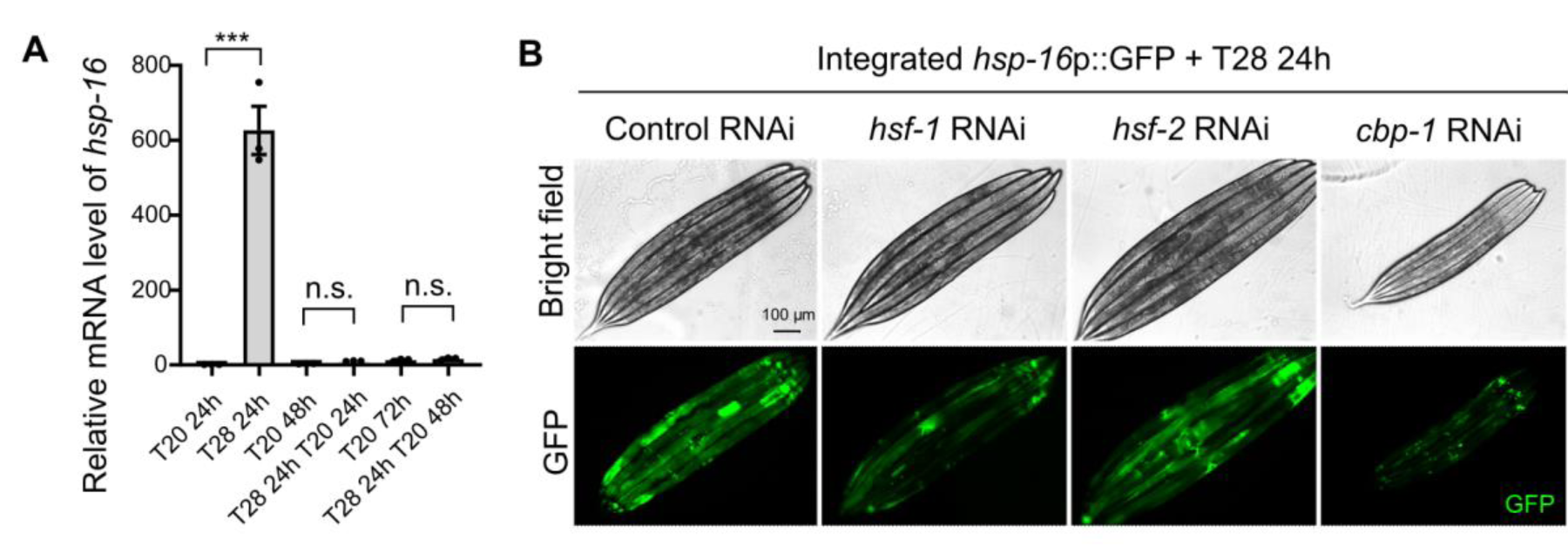
Heat shock protein induction by T28 requires both HSF-1 and CBP-1. (A) Quantitative RT-PCR measurements of *hsp-16* expression levels showing its transient induction by ELTS (T28 for 24 hrs at L4). *** indicates *P* < 0.001 (three independent biological replicates). (B) Representative epifluorescence images of animals with RNAi against *hsf-1* or *cbp-1* showing up-regulation of *hsp-16*p::GFP by ELTS that depends on both HSF-1 and CBP-1, but not HSF-2. Scale bars: 100 µm.

**fig. S7.**
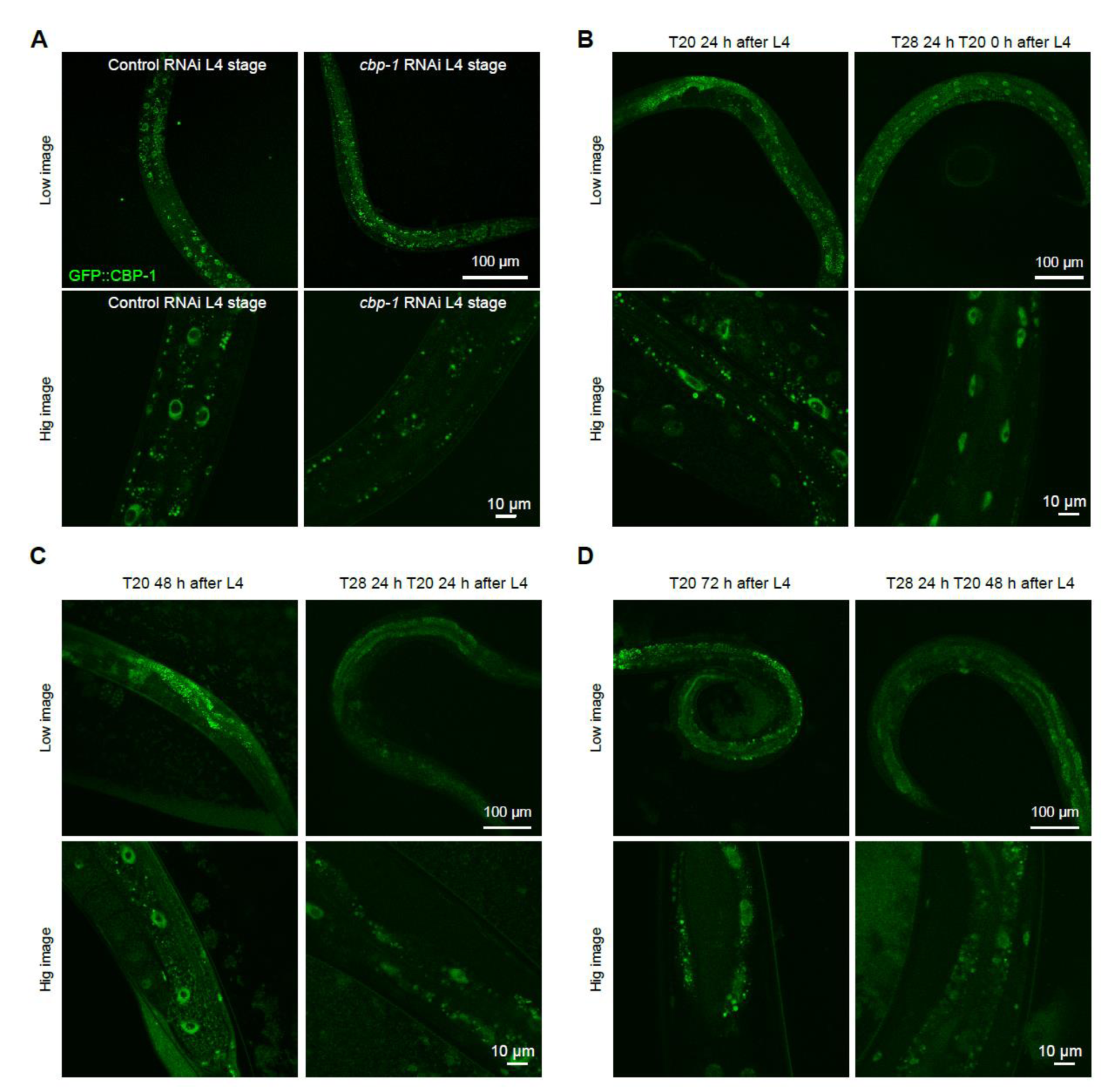
Heat-induced nuclear entry of CBP-1 endogenously tagged with GFP. (A) Representative confocal fluorescence images showing specific nuclear signals of GFP::CBP-1 (by CRISPR knock-in at the endogenous *cbp-1* locus) that were diminished by RNAi against *cbp-1*. (B) Representative confocal fluorescence images showing increased nuclear entry of endogenous GFP::CBP-1 by 28 °C for 24 hrs. (C) Representative confocal fluorescence images showing unaltered GFP::CBP-1 by 28 °C for 24 hrs and 20 °C for 24 hrs. (D) Representative confocal fluorescence images showing unaltered GFP::CBP-1 by 28 °C for 24 hrs and 20 °C for 48 hrs. Shown are both high and low-magnification views. Scale bars are indicated.

**fig S8.**
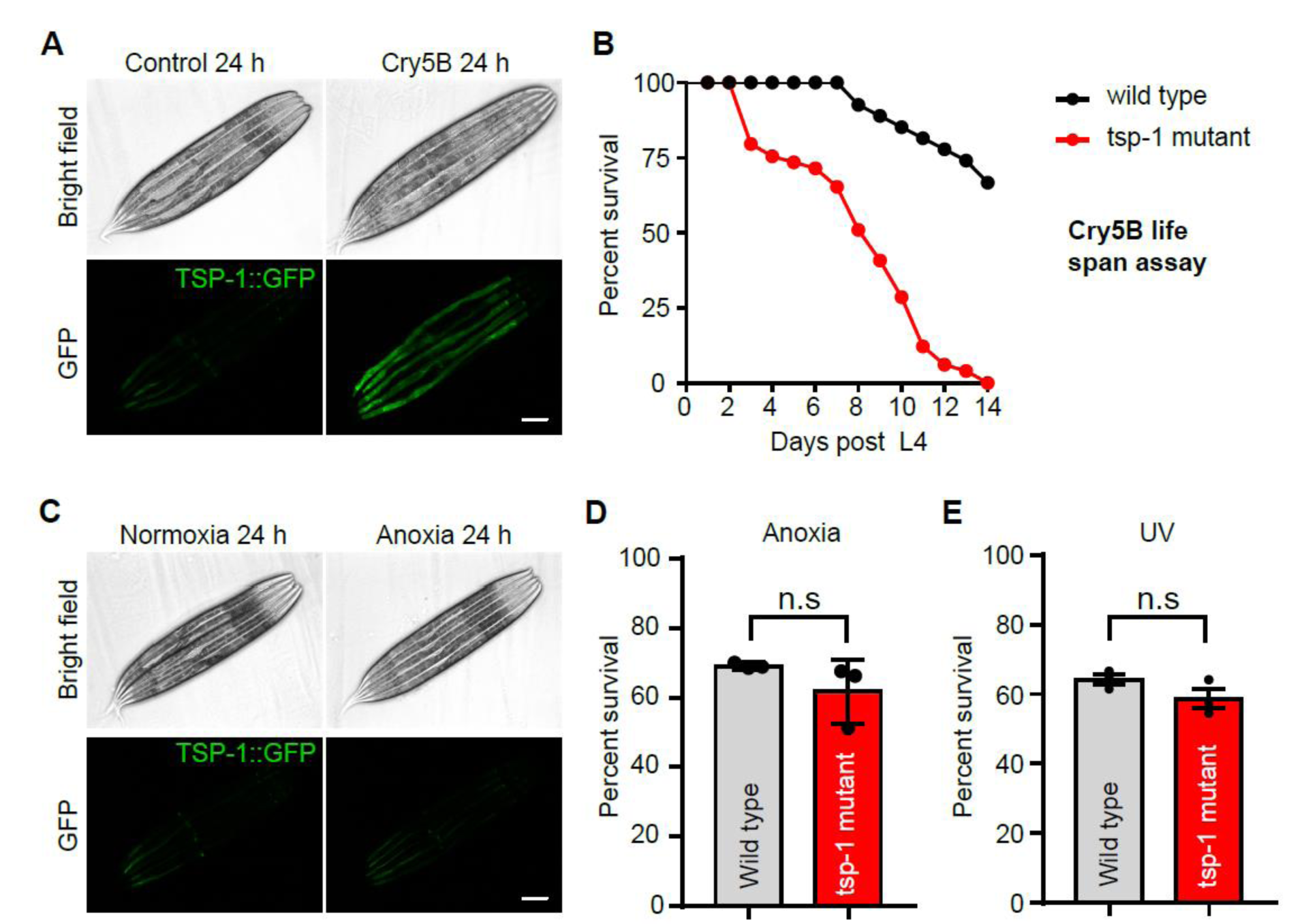
Cry5B, UV, and anoxia effects on TSP-1 and organismal stress resilience. (A) Representative brightfield and epifluorescence images showing expression of an integrated transgene *tsp-1*p::*tsp-1*::GFP treated with cry5B expressing bacteria or control. Scale bars: 100 μm. (B) Lifespan curves of wild type and *tsp-1* deletion mutant allele *ok3594* with cry5B or control starting at L4. (C) Representative bright field and epifluorescence images showing expression of an integrated transgene *tsp-1p::tsp-1*::GFP under normoxia or anoxia for 24 h. Scale bars: 100 μm. (D) Percentage survival of wild type and *tsp-1* deletion mutants after exposure to anoxia at L4. (E) Percentage survival of wild type and *tsp-1* deletion mutants after exposure to UV at L4.

**fig S9.**
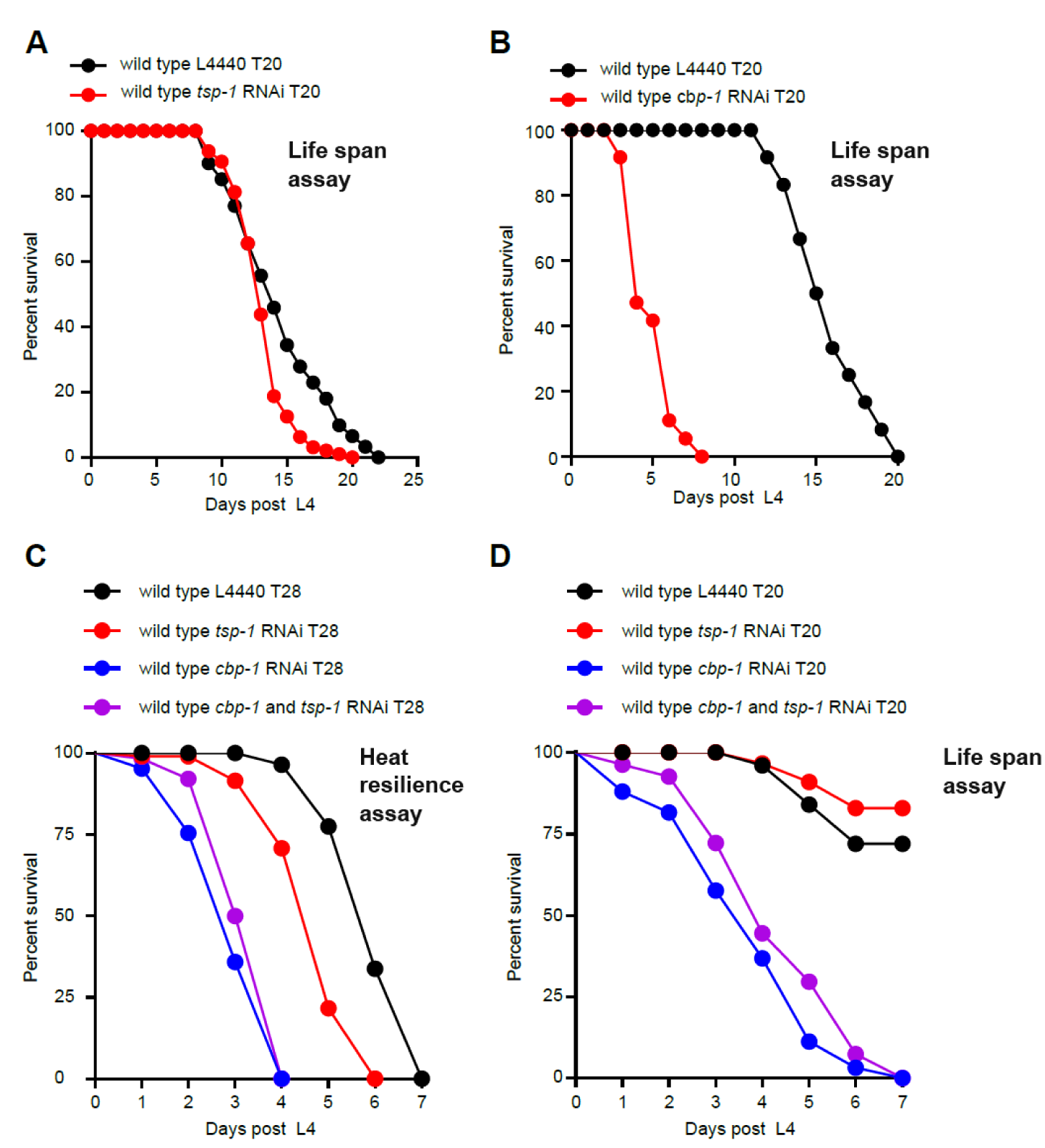
Effects of co-application of *cbp-1* RNAi and *tsp-1* RNAi in *C. elegans*. (A) Lifespan curves of wild type with *tsp-1* RNAi or control starting at L4. (B) Lifespan curves of wild type with *cbp-1* RNAi or control at constant 20 ℃. (C) Lifespan curves of wild type with *tsp-1* RNAi, *cbp-1* RNAi or *tsp-1*+ *cbp-1* RNAi exposure to 28 ℃ starting at L4. (D) Lifespan curves of wild type with *tsp-1* RNAi, *cbp-1* RNAi or *tsp-1*+ *cbp-1* RNAi at constant 20 ℃ starting at L4.

**Table S1.** Customized RNAi screen for candidate nuclear regulators of heat-induced TSP-1::GFP.

**Table S2.** Quantitative summary of lifespan assay statistics and results.

**Movie S1.** Tetraspanin webs exhibit stability from 120-min imaging of TSP-1::mScarlet. Scale bar: 10 µm.

